# Genetic approaches reveal a healthy population and an unexpectedly recent origin for an isolated desert spring fish

**DOI:** 10.1101/2023.07.07.548039

**Authors:** Brian L. Sidlauskas, Samarth Mathur, Hakan Aydoğan, Fred R. Monzyk, Andrew N. Black

## Abstract

Foskett Spring in Oregon’s desert harbors a historically threatened population of Western Speckled Dace (*Rhinichthys klamathensis*). Though recently delisted, the dace’s recruitment depends upon regular removal of encroaching vegetation. Previous studies assumed that Foskett Dace separated from others in the Warner Valley about 10,000 years ago, thereby framing an enigma about the population’s surprising ability to persist for so long in a tiny habitat easily overrun by plants. To investigate the phenomenon of persistence and the effectiveness of interventions to augment population size, we assessed genetic diversity among daces inhabiting Foskett Spring, a refuge at Dace Spring, and three nearby streams. Analysis revealed a robust effective population size (N_e_) of nearly 5,000 within Foskett Spring, though N_e_ in the Dace Spring refuge is just 10% of that value. Heterozygosity is slightly lower than expected based on random mating at all five sites, indicating mild inbreeding, but not at a level of concern. These results confirm the genetic health of Foskett Dace. Unexpectedly, genetic differentiation reveals closer similarity between Foskett Dace and a newly discovered population from Nevada’s Coleman Creek than between Foskett Dace and dace elsewhere in Oregon. Demographic modeling inferred Coleman Creek as the ancestral source of Foskett Dace just 600 years ago, much more recently than previously suspected and coincident with the arrival of large herbivores whose grazing may have maintained open water suitable for reproduction. These results solve the enigma of persistence by greatly shortening the duration over which Foskett Dace have inhabited their isolated spring.

## Introduction

The renowned ichthyologist Carl Hubbs opened a seminal work by observing pithily “where there is water, there are fishes” (Hubbs & Lagler, 1958). Indeed, fishes inhabit every imaginable aquatic habitat on our planet, from the deepest oceanic abyss (Gerringer et al., 2017) to barely moistened leaf litter (Britz et al., 2018), and from the freezing poles (Purser et al., 2022) to isolated pools in torrid deserts (Deacon & Minckley, 1974). Scientists studying these last, arid habitats often marvel at the ability of pupfishes, daces, gobies, salamanderfishes, mosquitofishes, rainbowfishes and others to persist in some of the world’s most marginal waters (Berra & Allen, 1989; Brown, 1971; Echelle et al., 1989; Mossop et al., 2023; Mussmann et al., 2020; Vinyard, 1996; Wager & Unmack, 2004). The surprising ubiquity of fishes in isolated desert systems invites exploration of their origins. How long ago did they reach their fragmented and distant homes, and by what means? How long have they endured there? Conservation biologists have also rightly raised concerns about the continued ability of these small populations of highly endemic species to persist in the face of changing landscape uses and a warming climate (Meffe & Vrijenhoek, 1988; Minckley & Deacon, 1991; Unmack & Minckley, 2008). And indeed, several of the earliest fishes listed as imperiled under the US Endangered Species Act inhabit small waters in the vastness of America’s arid west, such as the Devils Hole Pupfish (*Cyprinodon diabolis*), Desert Dace (*Eremichthys acros*), Owens River Pupfish (*Cyprinodon radiosus*) and Cui-ui (*Chasmistes cujus)* (Udall, 1967).

The origin, persistence and conservation of endemic snails, amphipods, plants and even planarians in desert springs has also attracted considerable attention (Inoue et al., 2020; Murphy et al., 2013; Murphy, King, et al., 2015; Robertson et al., 2014; Rossini et al., 2018; Rossini et al., 2020). These taxonomically disparate studies have confirmed that recent speciation and divergence contribute to the high endemism of desert waters (Campbell et al., 2022; Seidel et al., 2009; Witt et al., 2008), but also revealed that isolated springs can harbor relictual populations of ancient, formerly widespread lineages. For example, the isopod *Pheratomerus latipes*, a spring-dwelling endemic of the Lake Eyre region of South Australia, is a relict of a more broadly distributed lineage that diverged in the Miocene (Guzik et al., 2012). Stinking Lake Spring in Oregon’s high desert harbors a relictual lineage of Speckled Dace (*Rhinichthys osculus*) that diverged at least three million years ago from the daces that now inhabit the surrounding Malheur watershed and may merit recognition as a distinct species (Hoekzema & Sidlauskas, 2014). And indeed, some inhabitants of aquatic desert ecosystems radiated recently from within relictual lineages, making their spring habitats simultaneous cradles and museums of diversity (Murphy, Guzik, et al., 2015).

Given the risk of habitat alteration or the likelihood of small populations accumulating deleterious mutations through genetic drift, the persistence of lineages in desert springs over thousands to millions of years presents something of an enigma. Fervent debate has erupted over the age and origin of some iconic species, such as Nevada’s Devils Hole Pupfish, *Cyprinodon diabolis,* (Martin et al., 2016; Martin & Hohna, 2018; Sağlam et al., 2016), which maintains such a tiny population size that demographic models suggest a high probability of stochastic extinction over timescales exceeding a few thousand years (Martin et al., 2016; Reed & Stockwell, 2014). An exceptional ability to tolerate inbreeding and a relatively recent divergence time may explain how that pupfish has so far evaded extinction (Tian et al., 2022), but these factors may not generalize to other desert spring taxa. For some, such as the endemic desert spring amphipods of South Australia, relatively high genetic diversity and surprisingly large populations may insulate them from inbreeding and the deleterious effects of drift (Murphy et al., 2013). Occasional dispersal and gene flow augments the genetic diversity of other desert endemics such as *Fonscochlea* snails (Wilmer et al., 2008) *Chlamydogobius* fishes (Mossop et al., 2015) (Mossop et al., 2023) and *Ochthebius* beetles. (DeBoo et al., 2019). But, even in cases where occasional dispersal augments the genetic diversity of small populations trapped in tiny habitats, the ability of these taxa to avoid local extinction from stochastic alteration to their habitat remains remarkable.

This study investigates the genetic health and persistence of Foskett Spring Speckled Dace (Fig 1., hereafter Foskett Dace) an isolated Oregon population of *Rhinichthys klamathensis* that was listed federally as a threatened, undescribed subspecies of *R. osculus* between 1985 and 2019 (U.S. Fish and Wildlife Service, 1985, 2019a). The ichthyologist Carl Bond assigned probable subspecies status to Foskett Dace due to phenotypic differences from dace in the surrounding Warner Valley and because of its presumed isolation since the end of the late Pleistocene pluvial period (Carl Bond, Oregon State University, pers. comm. 1990, cited in U.S. Fish and Wildlife Service, 1998). Subsequent phylogeographic and population genetic studies discovered that Foskett Dace are identical or nearly identical to daces from elsewhere in the Warner Basin at maternally inherited mitochondrial loci (Ardren et al., 2009; Hoekzema & Sidlauskas, 2014) and concluded that they do not merit subspecies status alone. However, Foskett daces do differ enough in microsatellite allele frequencies to merit recognition as a distinct and significant population (Hoekzema & Sidlauskas, 2014) within a subspecies of Western Speckled Daces inhabiting the entire Warner Valley. Moyle et al. (2023) recognized this subspecies formally as *Rhinichthys klamathensis goyatoka*.

**Figure 1:**
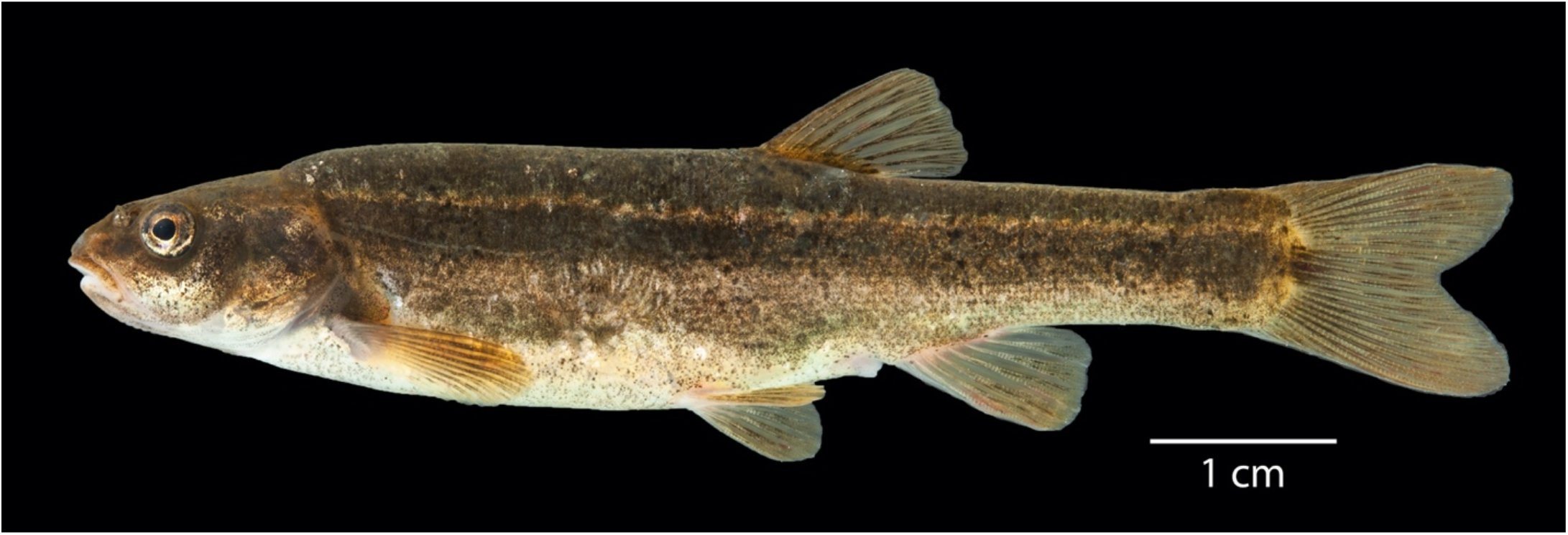
OS 22833, voucher BLS20-102, *Rhinichthys klamathensis goyatoka*, Foskett Spring, Oregon, photographed in an immersion tank immediately after euthanization.

Foskett Dace are endemic to a small, aquifer fed pool (Fig. 2, top) and an outflow stream that ends in a tule and cattail marsh on the shore of the normally dry Coleman lakebed (Scheerer & Jacobs, 2009). That spring lies within the Coleman subbasin of the Warner Basin, an endorheic basin located primarily in southeastern Oregon, though extending slightly into Nevada and California (Fig. 3). Previous studies (Ardren et al., 2009; Hoekzema & Sidlauskas, 2014) assumed that these fish came to occupy their tiny habitat about 10,000 to 12,000 years ago, during the final desiccation of pluvial Lake Warner (Hubbs et al., 1974; Wriston & Smith, 2017).

**Figure 2.**
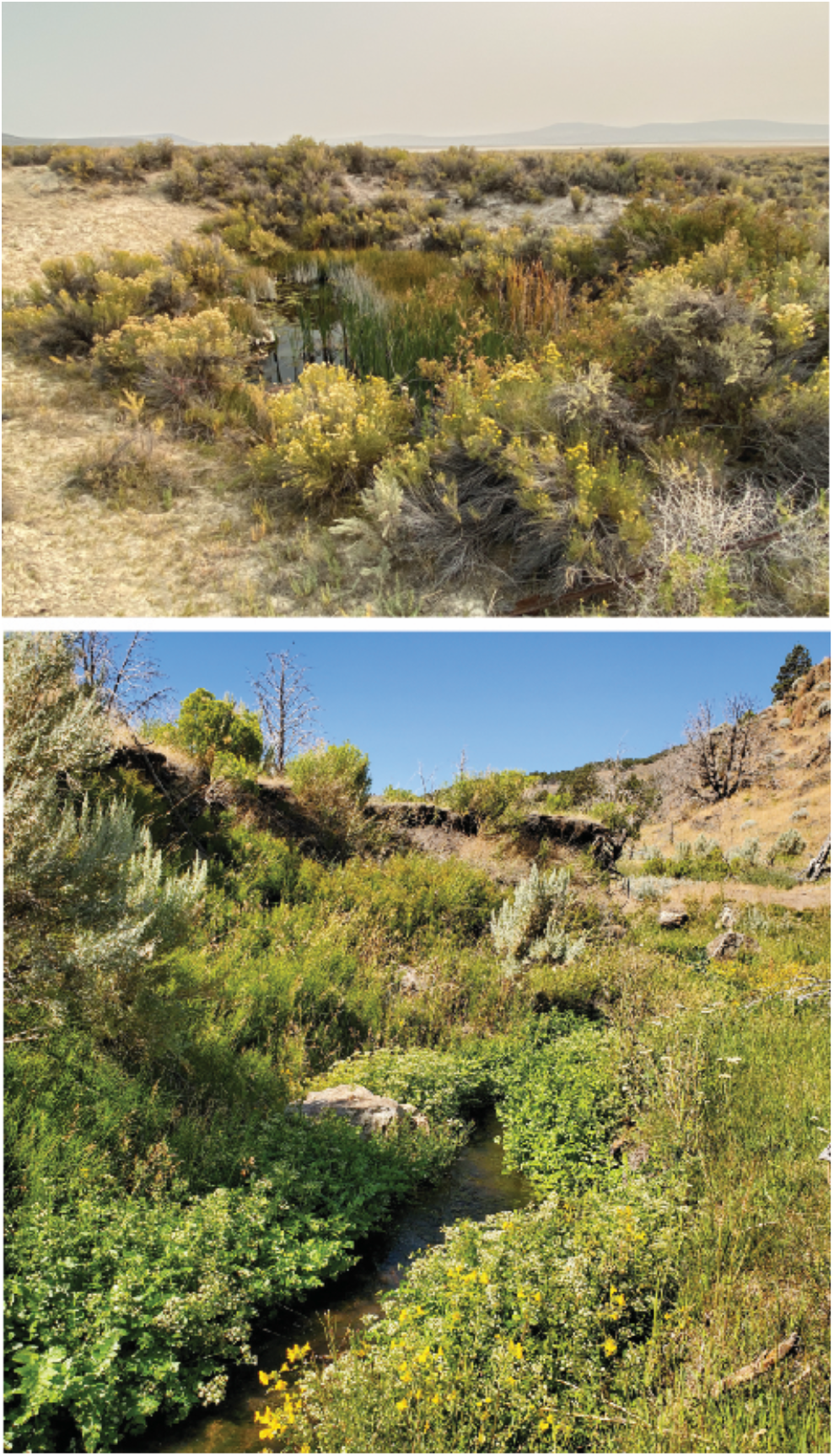
Collection sites for *Rhinichthys klamathensis goyatoka* at Foskett Spring, Oregon (top) and Coleman Creek, Nevada (bottom).

**Figure 3:**
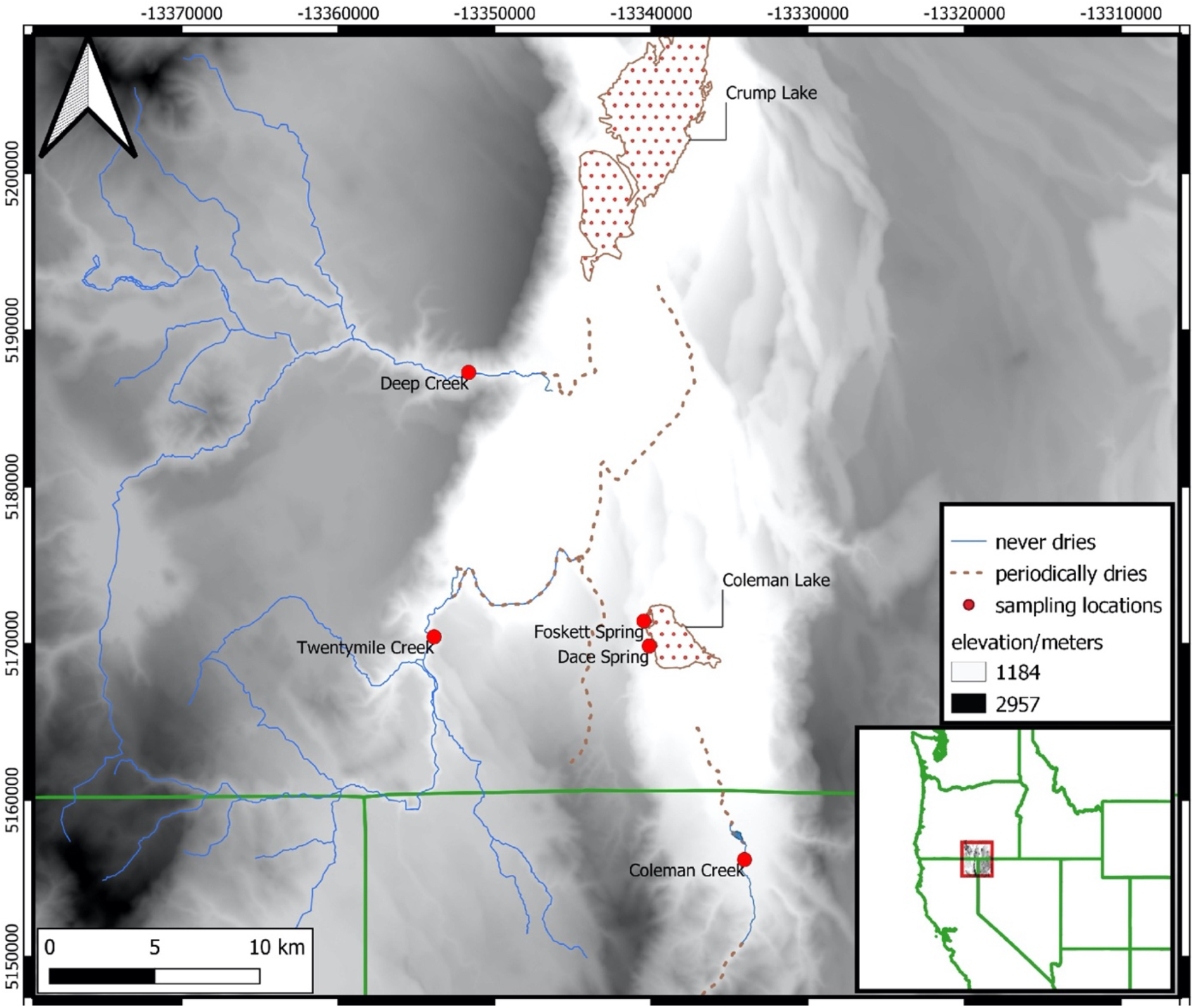
Map of sampling locations at and near Foskett Spring. Blue waters (solid lines) are wet year-round, while dashed regions indicate water bodies that fill only occasionally. Foskett Spring, Dace Spring and Coleman Creek lie within the Coleman subbasin of the Warner Valley. Deep and Twentymile creeks lie outside the Coleman subbasin. Inset map shows the location of the southern Warner Valley near Oregon’s borders with California and Nevada.

The population size of Foskett Dace fluctuates substantially from year to year, with the availability of open water habitat seeming to drive much of the variation. Presumably, the dace require open water for effective recruitment (U.S. Fish and Wildlife Service, 2009). Though historical data record a maximum of 27,787 individuals in 1997, from 2005 to 2011 estimates from a Lincoln-Peterson closed-capture estimator have averaged around 10% of that value. The decline coincides with a gradual disappearance of open water that began in 1987 when the Bureau of Land Management fenced the spring and its outflow system to exclude cattle (Scheerer et al., 2012). Census population size for Foskett Dace dropped to 1,728 individuals in 2011 during a period of extensive aquatic vegetation growth throughout the entire system, with abundances in 2012 and 2016 just slightly above that minimum (Peterson et al., 2015; Scheerer et al., 2012; Scheerer et al., 2017; assessments after 2011 used a Huggins closed-capture estimator). Population sizes rapidly increased in the years immediately following controlled burns or manual removal of aquatic vegetation to increase open water habitat, reaching a recent peak of 24,888 individuals in 2014 (Fig 4., Scheerer et al., 2017). Population sizes prior to the mid twentieth century are unknown, though the dependence of successful recruitment on the availability of open water habitat likely held.

**Figure 4:**
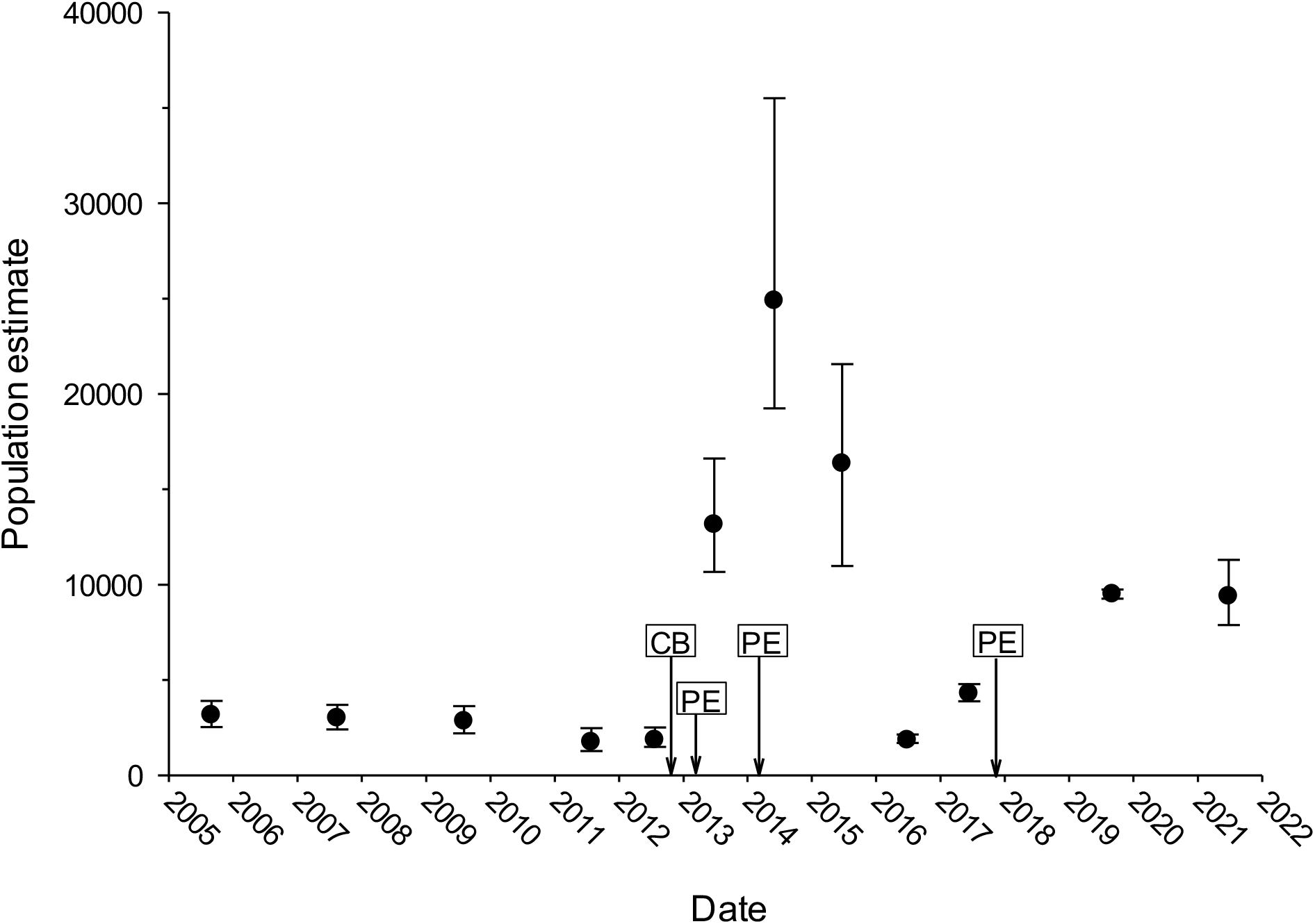
Foskett Spring Speckled Dace population abundance from 2005-2021. Arrows on the date axis indicate habitat enhancements to increase the open-water habitat throughout the spring system, and are offset by the year to indicate the month renovations occurred. CB=controlled burn, PE=pool excavation. Error bars are 95% confidence intervals. Data courtesy of the Oregon Department of Fish and Wildlife.

If Foskett Dace have indeed remained isolated throughout the dry conditions of the last 10,000 years, then their long persistence despite pronounced volatility in their population size raises an enigma. Though the capture-recapture studies did not calculate effective population sizes, N_e_ in wild populations is typically much lower than census population size, and sometimes only a tenth of that value (Frankham, 1995). Thus, if historical census population sizes for Foskett Dace have often fluctuated around a few thousand individuals, then effective population sizes have plausibly dipped below Franklin’s (1980) of-cited rule that N_e_ needs to remain above 500 to maintain the evolutionary potential of a population, and almost certainly below Frankham et al.’s (2014) revised threshold of N_e_ > 1000. The lowest census population sizes also fall short of Traill, Bradshaw, & Brook’s (2007) estimate for the minimum viable population size for animal species, which they calculated as 4,169 individuals through meta-analysis.

Given their tiny range, fluctuating population size, apparent dependence on open-water habitat for reproduction (U.S. Fish and Wildlife Service, 2009), and lack of an obvious migration pathway between Foskett Spring and the streams in the remainder of the Warner Valley for the last 10,000 to 12,000 years, how have they escaped extinction due to failed recruitment when vegetation overran their pool, or from the gradual erosion of genetic diversity? Does their population contain enough individuals of sufficient genetic diversity to insulate their lineage from effects of inbreeding and bottlenecks over thousands of generations? Are they lucky enough to have avoided failed recruitment despite encroaching vegetation? Did an unknown factor keep the spring pool clear? Or, is the population in Foskett Spring simply younger than suspected?

This contribution harnesses modern molecular methods to quantify the effective population size and levels of genetic diversity among Foskett Dace, the refuge population at Dace Spring, and daces in three nearby streams, including a newly discovered and never-before-sequenced population in Nevada’s Coleman Creek. By contributing a genetic assessment to demographic monitoring, these objectives help maintain continued health and viability of the recently delisted Foskett Dace. We also evaluate the discreteness of Foskett Dace relative to the other populations and infer their demographic history and date of divergence. In so doing, we reveal Coleman Creek as the true ancestral source of Foskett Dace and solve the enigma of their purported persistence for 10,000 years in a small spring system that modernly becomes quickly engulfed by vegetation without human intervention.

## Methods

### Study Area

The study examined specimens collected from five sites in two subbasins that connected hydrologically 12,000 years ago when Pleistocene Lake Warner filled the modern Warner Valley (Wriston & Smith, 2017). Foskett Spring, Dace Spring and Coleman Creek lie within the endorheic Coleman subbasin at the extreme southern end of the Warner Valley, while Twentymile and Deep Creeks lie within the main Warner subbasin to the north and west (Fig. 3). Though no current hydrographic connection exists between these subbasins, a low sill just one meter in height separates them at their point of closest contact, and the basin floors differ by only eight meters of elevation (Wriston & Smith, 2017). Though the modern Coleman subbasin does not connect hydrographically to the rest of the Warner basin, those small elevational differences imply that modest changes in water levels could re-establish connectivity.

Deep and Twentymile Creeks represent the major perennial streams of the southern portion of the main Warner Valley. Each terminates in shallow marshlands on the valley floor during low discharge but during periods of high discharge, waters from the creeks inundate large portions of the valley floor and flow northward through sloughs and irrigation canals into the semi-permanent Crump Lake (Hunt, 1964).

Within the Coleman subbasin, Foskett Spring (Fig 2, top) originates on the western slope of the normally dry Coleman Lake (Fig. 3), which fills only during rare heavy rainfalls (Williams et al., 1990), The spring forms a 33-m^2^ pool that feeds a small outflow channel for approximately 95 m and terminates in a shallow marsh at the edge of the dry lakebed. Adult and juvenile Foskett Dace inhabit all the wetted portions of this spring system. Dace Spring is a smaller spring system located just to the south of Foskett Spring. Though historically lacking fish, Dace spring was modified in 2010 to support a translocated refuge population of Foskett Dace. Across the Nevada border at the extreme southern end of the Coleman subbasin, Coleman Creek (Fig. 2, bottom) is a small intermittent stream perennially fed by small springs. Its short (∼5 km) reach terminates in an irrigation reservoir except during periods of high discharge when flows can reach Coleman Lake, implying the possibility of fish migration between Coleman Creek and Foskett Spring through Coleman Lake during unusually wet years. Prior to this study, the population of dace in Coleman Creek was unknown to science.

### Sample collection

Samples of adult daces that were used for genetic analyses were collected in 2020 from Coleman Creek (N=48), Foskett Spring (N=48), Dace Spring (N=48), Deep Creek (N=20) and Twentymile Creek (N=26) under Oregon Scientific Taking Permit 24050 and license 39884 issued by the Nevada Department of Wildlife. Fish were captured using minnow traps baited with sandwich bread and euthanized with tricaine methanesulfonate (MS - 222) at a concentration of 0.5g/L in natural spring water, in accordance with Animal Care and Use Protocol 2020-0115 approved by Oregon State University. Following euthanasia, selected specimens were photographed in an immersion tank (e.g., Fig. 1) and all specimens were immersed in ice water briefly while fin clips and plugs of epaxial muscle were collected. Voucher specimens for all tissue samples were collected and fixed in 10% formalin and have been accessioned into the Oregon State University Ichthyology Collection under catalog numbers OS22824, OS22836, OS22837, OS23148, OS22831, OS22832, OS22833, OS22834 and OS22835.

### Library preparation and sequencing

DNA extraction, library construction, and sequencing were all conducted at Oregon State University’s Center for Quantitative Life Sciences, largely following the methodology of Elshire et al. (2011). The main modification from this protocol involved the addition of a second restriction enzyme (see below). For each sample (N=190), genomic DNA was isolated from ∼50 mg of tissue using a DNeasy Blood and Tissue Kit (Qiagen), normalized to 20 ng/uL based upon Quibit fluorometer (Thermo Fisher Scientific) and fragmented using a double restriction digest with *pstI* and *mspI* on 200ng DNA. Following restriction enzyme digest, Illumina sequencing adapters and barcodes were ligated to individually fragmented DNA sequences. PCR products were cleaned using QIAquick PCR purification kit (Qiagen) and quality was assessed on an Agilent 2100 Bioanalyzer and loading concentration determined by qPCR. Pooled samples were then sequenced across two lanes of an Illumina HiSeq 3000, using paired-end 151-bp sequence chemistry.

### Quality filtering and SNP calling

Raw Illumina reads were assessed for quality using FASTQC (v0.11.9; Babraham Bioinformatics, Babraham Institute) and MULTIQC (v1.10; Ewels et al., 2016) prior to assembling sequence fragments into putative genetic loci using STACKS (v2.52; Catchen et al., 2013). The *process_radtags* script in STACKS was used to demultiplex raw sequences by assigning individual reads to corresponding samples through the unique combination of ligated in-line barcodes and standard Illumina read indices (Elshire et al., 2011). After removing the verified barcodes, truncated 141-base reads were checked for the presence of restriction cut-site sequences and only those reads containing correct motifs, or those with at most one base-call error (-r parameter), were retained. Using a sliding window of 21 bases (15% of read length), reads that showed an average decrease in quality score (Q score <10; 90% base-call accuracy) and those with any uncalled nucleotides were removed.

Demultiplexed reads were assembled into loci using the default parameters (but see below) of the STACKS *denovo_map.pl* wrapper program, which sequentially executes each core component of the pipeline. Retained loci were required to have a minimum allele depth of 5× (-m 5), be present in ≥80% of every population sample (--min-samples-per-pop 0.80) for all five populations (--min-populations 5) and have a minimum minor allele count of 3 (--mac 3). Furthermore, loci with an excessive number of reads (>2 SD above the mean depth) were filtered to remove any potentially merged paralogous loci. To maintain independence of loci with multiple polymorphisms, only the first SNP from each locus was retained. Furthermore, to screen close relatives in population samples, which could bias results (e.g., estimates of effective population size), KING-ROBUST (Manichaikul et al., 2010) was used to estimate the degree of relatedness between individuals using their kinship coefficient. The proposed cutoff values (Manichaikul et al., 2010) were used to identify and remove one individual from each pair of 1^st^ or 2^nd^ degree relatives discovered.

### Population genetic analyses

The filtered dataset was exported in PLINK file format (Purcell et al., 2007) and converted to various file formats using PGDSPIDER (v.2.0.5.2; Lischer & Excoffier, 2012). Effective population size (N_E_) was estimated using the linkage disequilibrium method implemented in the software package NEESTIMATOR (v.2.01; Do et al., 2014). GENODIVE (v.3.0; Meirmans, 2020) was used to calculate overall and per-locus observed heterozygosity (H_O_), expected heterozygosity (H_E_) and inbreeding coefficients (F_IS_). Overall and per-locus F_ST_ values (Weir & Cockerham, 1984) were calculated using DIVERSITY (v.0.04-22; Keenan et al., 2013) in R v.3.42 (R Core Team, 2019). To visualize groups of genetically similar individuals, a principal component analysis (PCA) was conducted on a matrix of allele frequencies using the package ADEGENET (v.2.01; Jombart, 2008) in R. For the PCA, data were scaled and centered, and missing values were replaced with mean population values.

Population structure was evaluated by estimating individual admixture proportions using the Bayesian clustering algorithm implemented in ADMIXTURE (v.1.30; Alexander et al., 2009). Model complexity was determined for each value of K (1–8) using the cross-validation function (-cv) to identify the value of K with the lowest associated error. The most likely model was then selected, and Q-scores were plotted with GGPLOT2 (v. 3.0; Wickham, 2016); for context, ±1K are plotted. We created a population-level neighbor-joining (NJ) dendrogram based on pairwise weighted *F_ST_* (Weir & Cockerham, 1984) differences between each population pair estimated using VCFTOOLS (Danecek et al., 2011).

To test for historic admixture between different populations and to identify the likely origin of the dace population in Foskett Spring, we used the three-sample D-statistic (*D_3_*) (Hahn & Hibbins, 2019). This D-statistic (also called the ABBA-BABA test) infers the post-divergence gene flow between divergent lineages based on gene genealogies (Green et al., 2010) and can test for gene flow of alleles among three populations with no outgroup (i.e., without polarization of alleles as ancestral or derived). Briefly, *D_3_* measures discordance in branch lengths in tree topologies based on pairwise genetic distances between different samples. If there are three samples A, B, and C where A is more closely related to B than to C, and *d_A-C_* is the genetic distance between A and C and *d_B-C_* is the genetic distance between B and C. Given those definitions, *D_3_* can be calculated as:

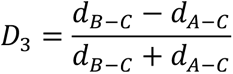

These discordance patterns should occur in equal frequencies between the two branches (A-C and B-C) and thus under expectation of no historic gene flow, *D_3_* = 0. Post-divergence gene flow between B and C would lead to *D_3_* > 0 and gene flow between A and C results in *D_3_* < 0. Since the *D_3_* statistic only requires a single sample from each population, we estimated genetic distances for each individual pair and estimated *D_3_* for all samples in each population trio in our dataset. We used *D_3_* estimates to identify the ancestral population of Foskett Dace by considering whether Coleman Creek or Deep Creek were the most likely source of gene flow into Foskett Spring, and therefore the probable ancestral population. These comparisons and the demographic model discussed below can only accommodate trios, not quartets, and so we chose Deep Creek as the exemplar locality from the population inhabiting the Warner Valley outside the Coleman subbasin (see results). Pairwise individual genetic distances were calculated using TASSEL (Bradbury et al., 2007) and *D_3_* for each population trio was calculated using R (R Core Team, 2019). We estimated the mean and 95% CI from the distribution of *D_3_* estimates for each population trio using the Z-distribution. We did not use the classic D-statistic because it estimates patterns of gene flow of a derived allele using an ancestral outgroup, and in this case the outgroup designation was uncertain because the order of colonization among the five localities was unknown.

### Reconstructing demography of Western Specked Dace populations

We reconstructed demographic history from the joint site frequency spectra (SFS) data with GADMA (v. 2.0.0rc20; Noskova et al., 2020) using the diffusion approximation as implemented in οaοI (Gutenkunst et al., 2009). This program estimates the population size (*N_e_*) changes, divergence times (*T*), and migration rates (*m*) among two or three populations. Given the limit of three populations, we modeled history using dace from Foskett Spring, Coleman Creek, and Deep Creek, each representing one of the three major populations identified by the PCA and Admixture analysis. Though Twentymile Creek dace are genetically similar to Deep Creek dace (see Results), they are not identical. We did not combine these subpopulations into a single population in the demographic models because this would have introduced metapopulation dynamics and subpopulation gene flow, which could render the results unreliable. We excluded the samples from Dace Spring because this is an artificial subpopulation with a known date of translocation.

For each demographic model, GADMA simulates the joint SFS and compares it to the observed SFS from population genetic data using log-likelihoods. It then chooses the most likely model by comparing AIC scores. To estimate confidence intervals (CI) of the parameter estimates (*N_e_, T, m*) for the most likely demographic model, we bootstrapped the observed joint SFS 100 times using independent subsets of SNPs. We converted genotype data to get observed SFS using EASYSFS (https://github.com/isaacovercast/easySFS) and for all demographic model simulations, we used a point mutation rate of µ=6.6×10^-8^, which is where Martin and Höhna (2018) centered the prior in their analysis of Devils Hole Pupfish., and which Recknagel et al. (Recknagel et al., 2013) measured as the actual mutation rate for RADSeq loci in the Midas cichlid (*Amphilophus* spp.). Speckled Dace rarely live for more than three years and never beyond four, and they typically reproduce at the age of two, though males may mature a year earlier than females (McPhail, 2007; Peden & Hughes, 1981; Wydoski & Whitney, 2003). Given the available information on life history, we set the generation time to two years, keeping all other parameters to default values.

## Results

### Quality filtering and SNP calling

Using 2×151 sequencing chemistry, the Illumina HiSeq 3000 yielded 758M reads across two sequencing lanes. Barcode sequences found within each multiplexed *fastq* file allowed assignation of paired-end reads to individual samples using the *process_radtags* script in STACKS. This resulted in 656M retained sequences (87.5%) after accounting for reads missing barcodes, restriction enzyme cut sites, or those exhibiting low base call quality. Following *de novo* assembly of loci among 190 individuals, samples had a mean depth of coverage (among 0.2M loci) of 134x. Among these genotyped loci, there remained 4,317 loci that met the sample (80%) / population (5) constraints. After removing two individuals that had a low genotyping rate (<70%) and eight individuals representing part of a pair of 1^st^ or 2^nd^ degree relatives, there remained 180 samples genotyped at 3,354 SNPs.

### Population genetic analyses

Using the linkage disequilibrium method, effective population sizes were moderately low for Dace Spring (N_E_=506), high for Coleman Creek (N_E_=1,418) and Foskett Spring (N_E_=4,950) and infinite for the Deep and Twentymile creek populations (Table 1). To assess the genetic health of the populations, we calculated overall and per-locus observed heterozygosity (H_O_), expected heterozygosity (H_E_) and inbreeding coefficients (F_IS_). All five populations showed similar levels of H_O_ (0.156-0.171) while the creek populations (0.175-0.181) exhibited marginally higher levels of H_E_ than the spring populations (0.163-0.164; Table 2). The close relationship between expected and observed diversity resulted in very low F_IS_ values among all Western Speckled Dace populations (0.036-0.057; Table 2). Pairwise F_ST_ estimates (Table 3) illustrated that Foskett Spring and Dace Spring daces have highly similar gene pools (F_ST_=0.0055), which is not surprising given that Dace Spring lacked fish until ∼100 daces were transplanted there from Foskett Spring in 2010 and 2011. Twentymile Creek daces differ moderately from those inhabiting Deep Creek (F_ST_=0.0604). The comparisons involving Coleman Creek showed moderate genetic differentiation from the two spring populations (F_ST_=0.0671-0.0704), and twice that difference from the two creek populations outside of the Coleman subbasin (F_ST_=0.1128-0.1275). Thus, daces in Foskett Spring are genetically more like Coleman Creek daces than they are like Twentymile or Deep Creek daces (F_ST_=0.1467-0.1605).

**Table 1:**
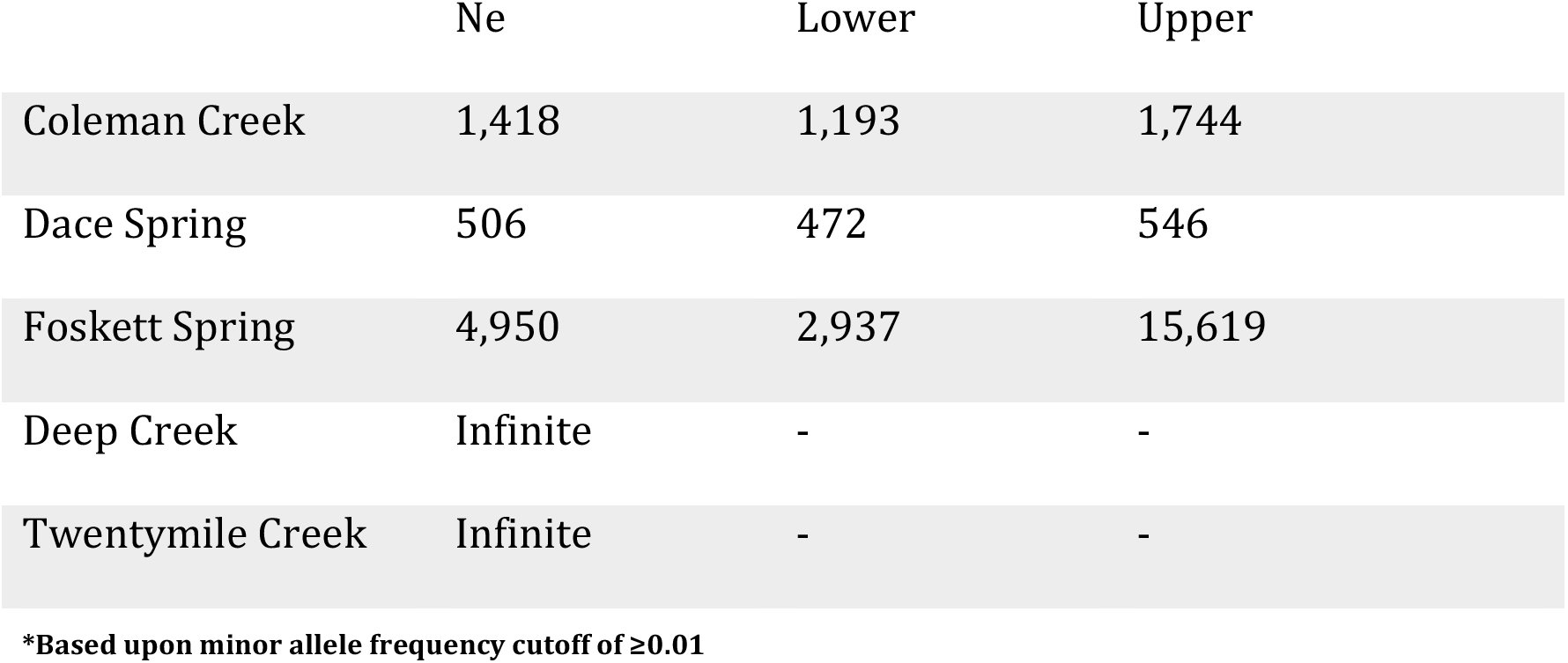
Effective population size estimates for the five sampled populations, calculated using the linkage disequilibrium method as implemented in NeEstimator (V.2).

|  | Ne | Lower | Upper |
| --- | --- | --- | --- |
| Coleman Creek | 1,418 | 1,193 | 1,744 |
| Dace Spring | 506 | 472 | 546 |
| Foskett Spring | 4,950 | 2,937 | 15,619 |
| Deep Creek | Infinite | - | - |
| Twentymile Creek | Infinite | - | - |

\*Based upon minor allele frequency cutoff of $\geq 0.01$

**Table 2:**
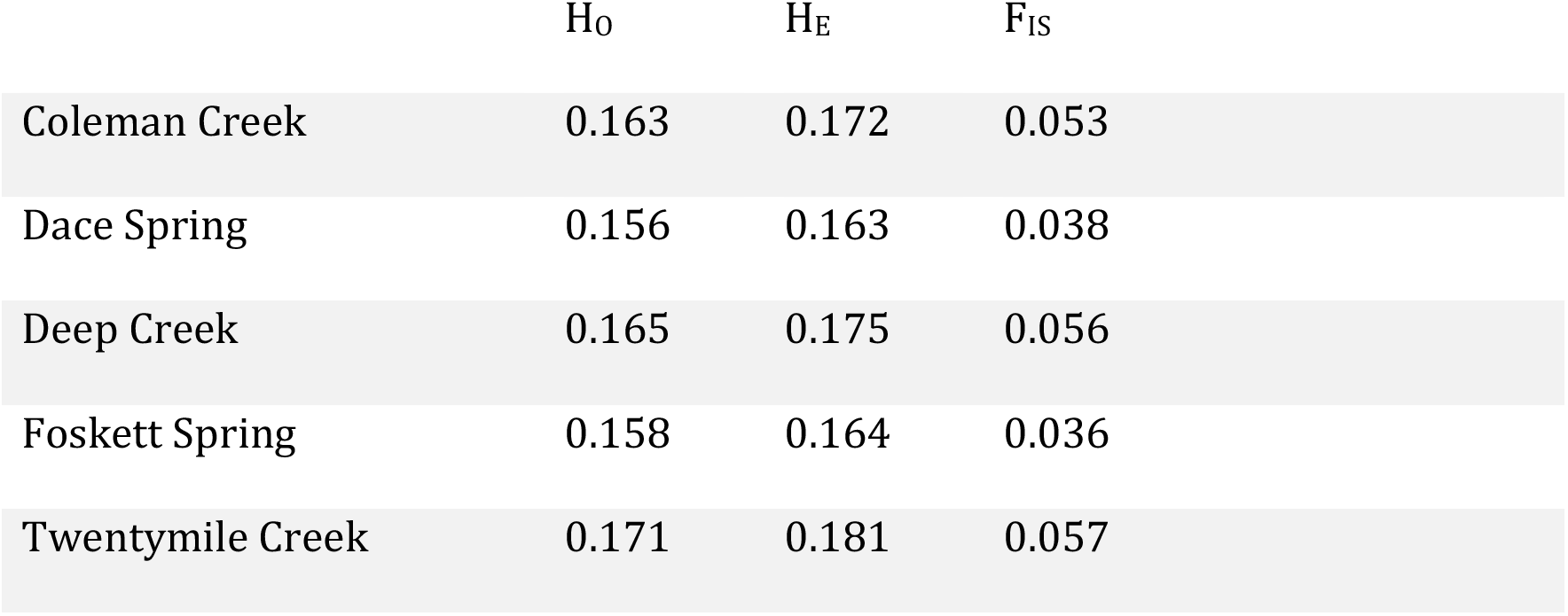
Population genetic metrics for the five Western Speckled Dace populations prior to removing relatives. Mean observed (H_O_) and expected (H_E_) heterozygosity, and mean inbreeding coefficient (F_IS_).

**Table 3:** Pairwise global F_ST_ values among the five sampled Western Speckled Dace populations.

|  | Coleman Creek | Dace Spring | Deep Creek | Foskett Spring |
| --- | --- | --- | --- | --- |
| Coleman Creek |  |  |  |  |
| Dace Spring | 0.0704 |  |  |  |
| Deep Creek | 0.1275 | 0.1644 |  |  |
| Foskett Spring | 0.0671 | 0.0055 | 0.1605 |  |
| Twentymile Creek | 0.1128 | 0.1499 | 0.0604 | 0.1467 |

PCA of allele frequencies clustered the individuals from Foskett and Dace springs together, united individuals from Twentymile and Deep Creek, and separated Coleman Creek daces cleanly from the other clusters (Fig. 5). PC1 (explaining 9.4% of variance) separates all three clusters, though it places Coleman Creek daces closer to the individuals from the spring populations than the other creeks. PC2 (3.6% of variance) clearly captured the variation separating Coleman Creek from all other populations (Fig. 5). Cross validation of K 2-8 identified K=3 as the having the lowest associated error. This model clearly clustered the two spring populations (Foskett and Dace) together, placed Coleman Creek dace in their own cluster, and grouped dace from Deep and Twentymile creeks (Fig. 6A). While associated error for the hypothesis of K=4 exceeded that for K=3, the clean separation of dace from Deep and Twentymile creeks under the four-group model implies subpopulation-level divergence between those two sites in the main Warner basin. Samples from Coleman Creek separated clearly from the other groups assuming K=3 or 4, albeit with low to moderate admixture with the spring samples for K=2-4 (Fig. 6A). Coleman Creek is the only population to return a dominant signal of admixture at K=2. A neighbor joining dendrogram constructed from a weighted *F_ST_* matrix of all pairwise populations concurs with the intermediacy of Coleman Creek dace relative to the Foskett and Dace springs fish on one hand, and the fish from Twentymile and Deep creeks on the other (Fig. 6C).

**Figure 5.**
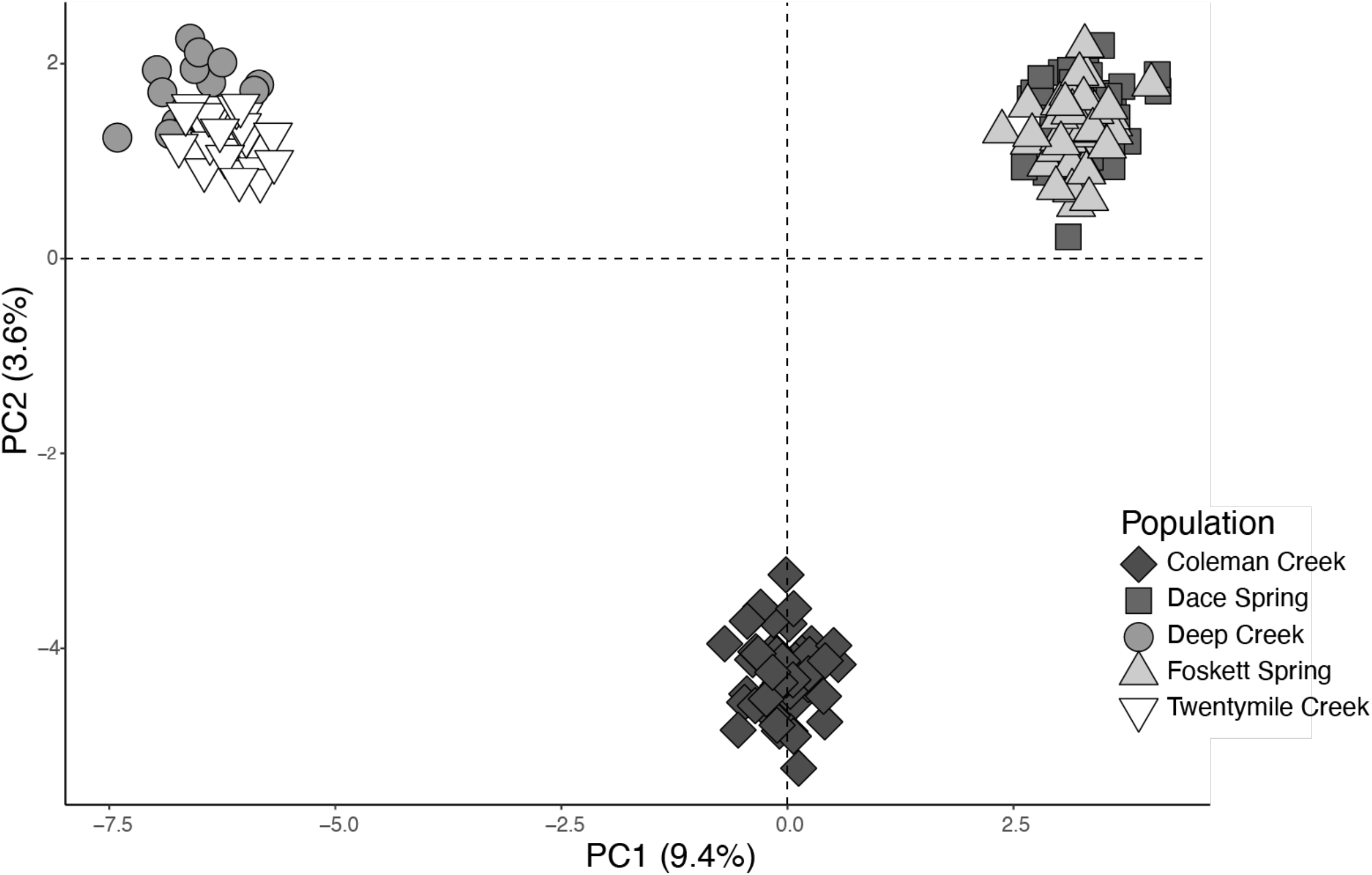
Principal Component Analysis of Western Speckled Dace in the southern Warner Valley. First two axes (PC1 & PC2) generated from allele frequencies among the five sampled dace populations

**Figure 6.**
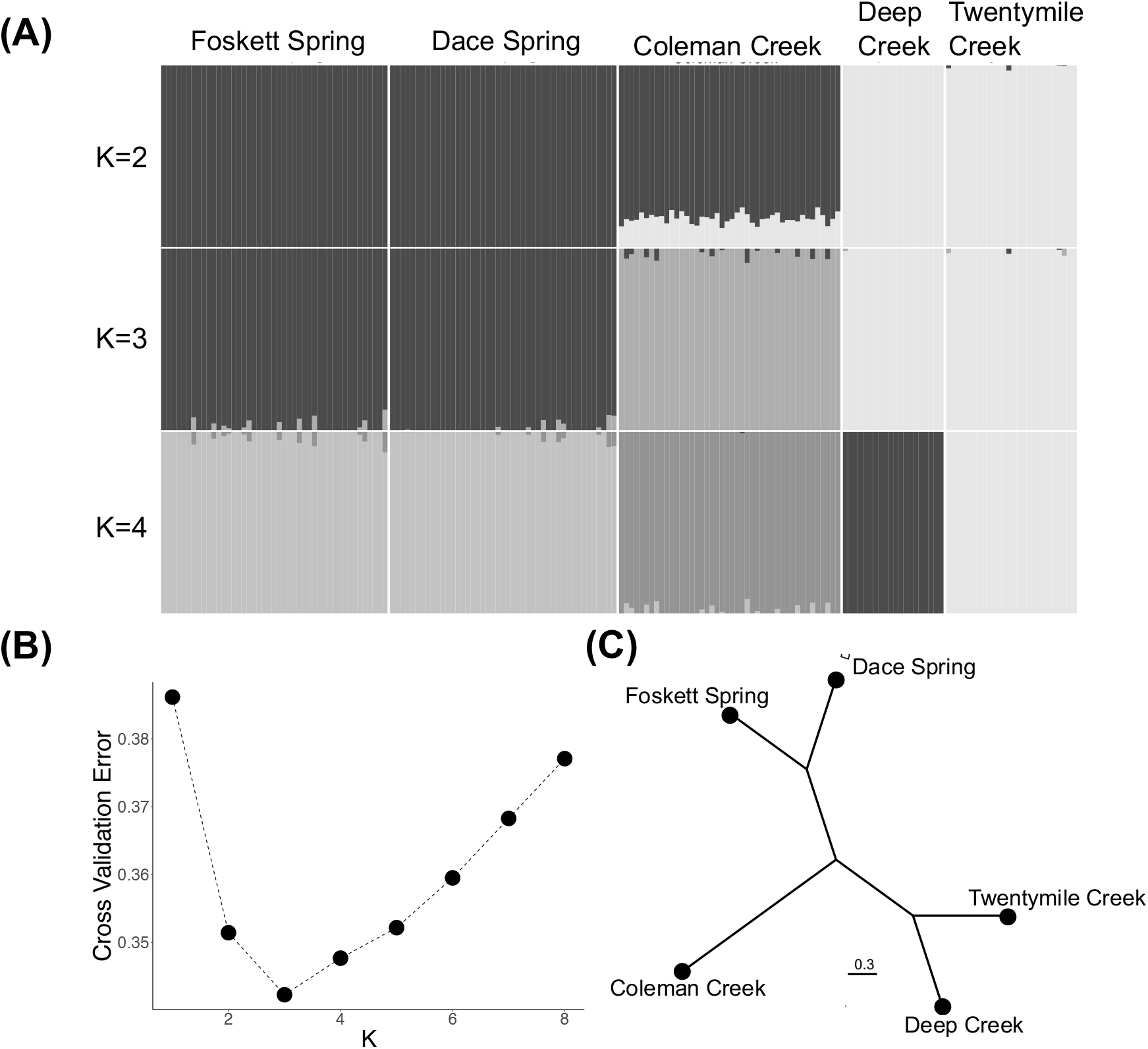
Population structure of Western Speckled Dace in the southern Warner Valley. **(A)** Population structure plot showing ancestry proportions for each individual (Coleman Creek = 44, Dace Spring = 45, Deep Creek = 20, Foskett Spring = 45, Twentymile Creek = 26) at different numbers of ancestral populations (*K*). The model identified K=3 as the most likely number of ancestral populations **(B)** Cross validation errors for all tested values of K in the population structure analysis shown in panel A. **(C)** Neighbor joining dendrogram constructed from a weighted *Fst* matrix of all pairwise populations.

To explore historic admixture (i.e., gene flow) among the three geographically and genetically separate populations, we estimated the *D_3_* -statistic for a trio of localities (Foskett Spring, Coleman Creek, Deep Creek) representing the three major genetic clusters. We considered scenarios with either (a) Foskett Spring and Deep Creek most closely related, or (b) Foskett Spring and Coleman Creek most closely related. If the first scenario is correct, we would expect no gene flow between Coleman Creek and Foskett Dace (*D_3_* = 0), yet we uncover strong evidence of gene flow between the Coleman Creek and Foskett Dace populations (*D_3_* = 0.054; 95% CI = [0.0527-0.0558]) (Fig. 7A). This suggests that the scenario of an initial split separating the Coleman Creek population from a lineage leading to the Foskett and Deep Creek populations is incorrect. On the other hand, D3 values more closely match predictions if we consider scenarios in which Deep Spring is ancestral and the Foskett Spring and Coleman Creek populations diverged more recently (Fig. 7B-C). In a scenario that considers Deep Creek to be most ancestral and postulates that Foskett Spring dace diverged from the Coleman Creek population (Fig. 7B), we find no evidence of historic gene flow between Foskett Spring and Deep Creek populations and weak evidence of historic post-divergence gene flow between Coleman Creek and Deep Creek populations (*D_3_* = -0.005; 95% CI = [-0.0066, -0.0033]). Similar weak evidence of historic gene flow between Coleman Creek and Deep Creek obtains if we consider Foskett Spring to be ancestral to the Coleman Creek population (Fig. 7C; *D_3_* = 0.0046; 95% CI = [0.0030, 0.0062]). Taken together, these results indicate that the deepest split occurred between the Deep Creek population and the lineage leading to the Coleman Creek and Foskett Spring populations. Historic gene flow between Coleman Creek and Deep Creek may then have occurred after the split between the Coleman Creek and Foskett Spring populations.

**Figure 7:**
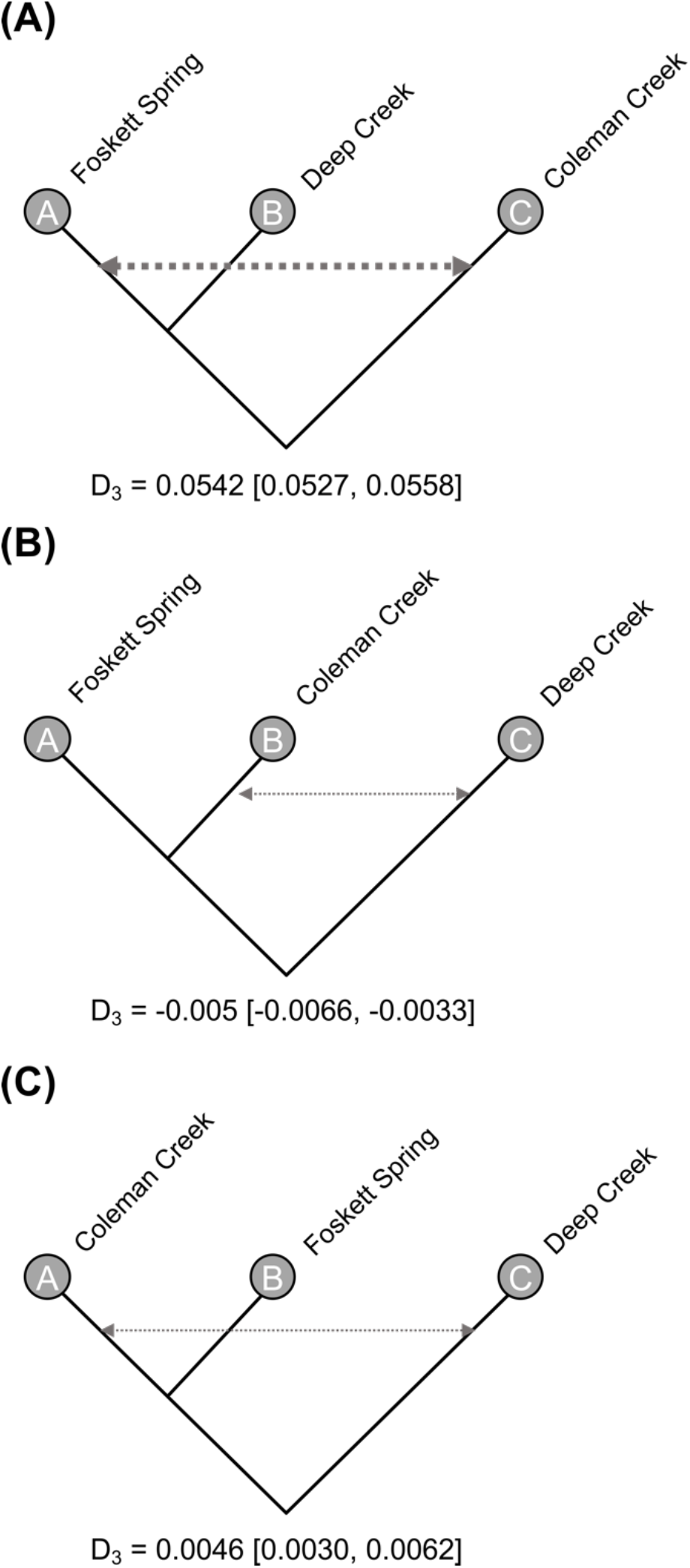
Inference of historic admixture between Western Speckled Dace populations. We tested for evidence of historic gene flow between diverging populations by measuring the *D_3_*-statistic among population trios. In each case, population A is more closely related to B than to C. D3 measures the discordance in branch lengths with *D_3_* = 0 means no gene flow, *D_3_* > 0 indicates gene flow between A and C, whereas *D_3_* < 0 represents gene flow between B and C. We estimated *D_3_* statistics for tree topologies where **(A)** Coleman Creek was sister to Foskett Spring and Deep Creek and **(B-C)** Deep Creek was sister to Foskett Spring and Coleman Creek. We find strong evidence of gene flow (shown by arrows) between Foskett Dace and Coleman Creek (A) and weak evidence of gene flow between Coleman Creek and Deep Creek (B-C). Numbers below each figure indicate mean *D_3_* for the given topology with 95% CI around the mean given in brackets. Thickness of arrows represent strength of gene flow. These results indicate that Foskett Dace likely diverged most recently from the Coleman Creek population, and that the Coleman Creek population experienced post-divergence gene flow with Deep Creek, possibly via secondary contact.

### The ancestors of Coleman Creek dace founded the Foskett Spring population

We estimated divergence times and long-term N_e_ from a three-population demographic model constructed by simulating joint site frequency spectra of the Western Speckled Daces inhabiting three localities (Coleman Creek, Deep Creek, Foskett Spring), representing the three modern populations. Table 4 lists the parameter estimates. The most likely demographic model indicates that an ancestral population of just under 2000 individuals (*N_anc_* = 1,890; 95% CI = [1,789 – 1,923]) split into two populations (P_11_ and P_12_) about 4000 years ago [best estimate of 3,784 years, 95% CI of 3,712 – 3,878] with 46.5% of the ancestral population forming P_11_ in the main Warner Valley and 53.5% of the ancestors forming P_12_ in the Coleman subbasin (Fig. 8). After that initial split, the lineage in the main Warner basin (P_11_) grew linearly to become effectively larger than Coleman Valley population (P_11_) (*N_p11_* = 5,863; *N_p12_* =1,011) until an event about 588 years ago [95% CI of 572 – 595] that restructured both lineages. The P_11_ population (leading to the modern Deep Creek subpopulation of the population in the main Warner Valley, and presumably also the subpopulation in Twentymile Creek) underwent a strong bottleneck, while P_12_ split into the Coleman Creek population and Foskett Spring population, with 88.7% of these P_12_ ancestors forming the Coleman Creek population. These results indicate that the modern diversity of Western Speckled Dace in the Warner Valley derived from two ancestral lineages, each confined to a different subbasin, and each affected substantially by events occurring 600 years (∼300 generations) ago.

**Figure 8:**
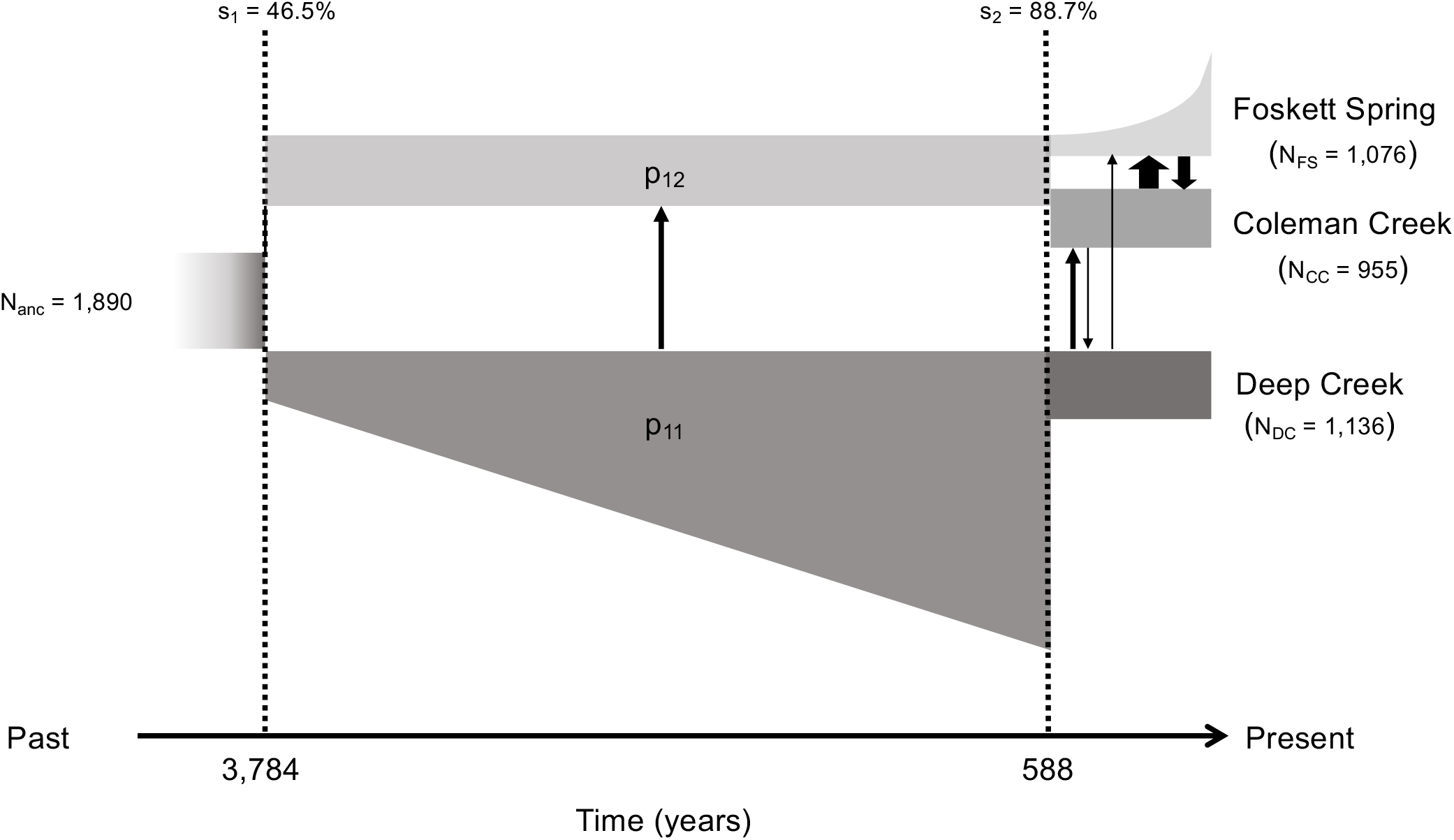
The most likely three-population demographic model for Western Speckled Dace in the southern Warner Valley, constructed by simulating joint site frequency spectrum data using GADMA with µ=6.6×10^-8^ and a generation time of 2 years. Mean N_e_ and divergence time estimates are presented here; all parameters and 95% confidence intervals around mean estimates are provided in Table 4. Arrows represent direction of post-divergence migration (gene flow) among the populations, with thickness indicating the magnitude of the migration rate. Bison arrived in the southern Warner Valley between 642 to 257 years ago, roughly coincident with the demographic shifts in dace populations inferred at 588 years before present.

**Table 4:** Maximum likelihood parameters for three-population demographic model for Western Speckled Dace inhabiting Coleman Creek, Deep Creek, and Foskett Spring inferred using GADMA. Values within parenthesis represent 95% confidence intervals estimated by bootstrapping observed site frequency spectra 100 times.

|  |  |
| --- | --- |
| No. of parameters estimated | 17 |
| Log likelihood | -9239.14 |
| Population size [95% CI] |  |
| $NA_0$ | 1,890 [1,789 – 1,923] |
| $P11_0$ | 879 [866 – 884] |
| $P11_1$ | 5,863 [5,813 – 5,901] |
| $P12_0 (= P12_1)$ | 1,011 [998 – 1,094] |
| $FS_0$ | 114 [105 – 125] |
| $FS_1$ | 1,076 [1,023 – 1,088] |
| $DC_0 (= DC_1)$ | 1,136 [1,112 – 1,152] |
| $CC_0 (= CC_1)$ | 955 [934 – 975] |
| Migration rate per generation (95% CI) |  |
| $m_{p11-p12}$ | $1.09 \times 10^{-2}$ [$1.03 \times 10^{-2}$ – $1.12 \times 10^{-2}$] |
| $m_{DC-CC}$ | $8.04 \times 10^{-3}$ [$8.03 \times 10^{-3}$ – $8.05 \times 10^{-3}$] |
| $m_{CC-DC}$ | $4.00 \times 10^{-3}$ [$3.97 \times 10^{-3}$ – $4.02 \times 10^{-3}$] |
| $m_{DC-FS}$ | $3.08 \times 10^{-3}$ [$3.04 \times 10^{-3}$ – $3.09 \times 10^{-3}$] |
| $m_{FS-DC}$ | 0 |
| $m_{CC-FS}$ | $2.65 \times 10^{-2}$ [$2.61 \times 10^{-2}$ – $2.68 \times 10^{-2}$] |
| $m_{FS-CC}$ | $1.61 \times 10^{-2}$ [$1.57 \times 10^{-2}$ – $1.64 \times 10^{-2}$] |
| Divergence time in years (95 % CI) |  |
| $T_1$ | 3,783.8 [ 3,712.4 – 3,877.9] |
| $T_2$ | 587.5 [ 572.4 – 595.1] |

*NA\_0: size of ancestral population; P11\_0: size of P11 immediately after divergence from* *ancestral population; P11\_1: size of P11 after linear expansion; P12\_0: size of P12* *immediately after divergence from ancestral population; FS\_0: size of Foskett Spring* *population immediately after divergence from P12; FS\_1: size of Foskett Spring population* *after exponential expansion; DC\_0: size of Deep Creek population immediately after* *bottleneck from P11; CC\_0: size of Coleman Creek population after divergence from P12; $m_{ab}$ :* *migration rate from population a to b; T\_1: divergence time between P11 and P12; T\_2:* *divergence time between Foskett Dace and Coleman Creek population.*

The dynamics of the three modern populations have differed over the most recent centuries. Though it was founded by only 11.3% of the Coleman valley ancestors, the contemporary Foskett Spring population grew exponentially to its current sizes (*N_FS_* = 1,076 [1,023 – 1,088]), whereas the contemporary Coleman Creek population remained stable after separating from the ancestral population, other than for experiencing a weak bottleneck at the time of separation (*N_CC_* = 955 [934 – 975]). Since the historic event that reduced effective population size circa 588 years ago, the contemporary Deep Creek subpopulation has remained stable (*N_DC_* = 1,136 [1,112 – 1,152]).

Our model inferred gene flow occurring at several points over the last few millennia. Unidirectional historic gene flow from the main Warner basin to the Coleman subbasin l populations (*m_p11-p12_* = 1.09×10^-2^) occurred between their initial subdivision 4000 years ago and the event that founded the Foskett population about 600 years ago. More recently, the demographic model inferred unequal and unidirectional gene flow among the three contemporary populations. Gene flow from Deep Creek into both Coleman Creek and Foskett Spring was identified with higher migration rate from Deep Creek to Coleman Creek (*m_DC-CC_* = 8.04×10^-3^) than from Deep Creek to Foskett Spring (*m_DC-FS_* = 3.08×10^-3^). Low levels of gene flow from Coleman Creek to Deep Creek were also identified (*m_CC-DC_* = 4.00×10^-3^) but the demographic model estimated no gene flow from Foskett Spring to Deep Creek (*m_FS-DC_* = 0). The highest migration rate was identified between Coleman Creek and Foskett Spring with more gene flow from Coleman Creek to Foskett Spring (*m_CC-FS_* = 2.65×10^-2^) than from Foskett Spring to Coleman Creek (*m_FS-CC_* = 1.61×10^-2^).

## Discussion

### Three distinct populations of Western Speckled Dace inhabit the southern Warner Valley

All our population genetic results clearly indicate three distinct clusters of Western Speckled Dace, each confined to a distinct hydrographic system within the southern Warner Valley (Figs. 5,6A). Analysis of genome-wide SNPs from GBS readily distinguished Foskett Spring dace from conspecifics collected from Twentymile or Deep Creeks, confirming the results from an earlier microsatellite study (Hoekzema & Sidlauskas, 2014). Fish from the newly discovered locality in Coleman Creek form a third cluster that was readily distinguishable from the other groups in all analyses. As expected, given their origin as a recent translocation, fish in Dace Spring are not distinguishable from those in Foskett Spring.

Fish from the two stream localities outside the Coleman subbasin cluster separately in neighbor joining dendrograms and in an Admixture analysis with K=4 (Fig. 6). However, K=4 is only the second most likely model, and the most likely model (K=3) groups dace from Twentymile and Deep Creeks into a single population. We interpret these patterns as reflecting separate, recently diverged subpopulations within the population that occupies the Warner Valley outside of the Coleman subbasin. Wiesenfeld et al. (2018) obtained similar results from Western Speckled Dace within California’s Klamath-Trinity basin, where they discovered three genetic clusters despite the lack of any obvious physical barrier separating two of them. If this kind of substructure within river systems is typical for *Rhinichthys klamathensis*, it might indicate that individuals tend towards strong site fidelity. Alternatively, sampling at more sites within the Twentymile and Deep stream networks might reveal intermediate individuals that blur the genetic distinction.

Overall, these results parallel other recent studies of *Rhinichthys* by recovering a genetically distinct population in each isolated drainage (Mussmann et al., 2020; Su et al., 2022). Similar results have been obtained in many other fishes inhabiting desert springs (Echelle et al., 1989; Faulks et al., 2017), dendritic riverscapes (Hopken et al., 2013; Morrissey & de Kerckhove, 2009; Osborne et al., 2014), or both (Echelle et al., 1987). Both situations create landscapes that can restrict gene flow, either by stranding fishes in isolated pools and springs, or by separating headwater populations by many river miles (Meffe & Vrijenhoek, 1988). At the same time, it is worth noting that not every desert spring necessarily harbors fishes representing a distinct population. For example, Campbell et al. (2022) found just three distinct genetic clusters among ten relictual spring localities in their study of *Crenichthys baileyi* in southeastern Nevada. Thus, while spring populations do often represent important components of genetic diversity, their distinctiveness must be tested as Stockwell et al (2013) did in their study of spring-dwelling populations of *Cyprinodon tularosa*, not assumed.

### Foskett Dace are genetically healthy and stable

Despite their isolation and tiny habitat, Foskett Dace appear to be doing remarkably well. We estimated their effective population size at 4,950 fish, which triples the value for the probable source population in Coleman Creek (Table 1), exceeds Frankham et al.’s (2014) threshold of N_e_ > 1000 to maintain population viability over evolutionary time, and surpasses the median value for minimum viable population size in animals (4,169) calculated in a meta-analysis of three decades worth of case studies (Traill et al., 2007). The large N_e_ for Foskett Spring based on the samples collected in 2020 reconciles well with the 2019 demographic estimate of 7,354 (95% CI 6,975–7,833) ‘adult-sized’ dace (fish > 35mm fork length) using a mark-recapture method (U.S. Fish and Wildlife Service, 2019a), particularly given that Speckled Dace do not begin to reproduce until they reach around 60 mm (McPhail, 2007). These numbers are substantially higher than some historical estimates, which have dipped below 2000 individuals in years of vegetation encroachment, as they did in 2011 (Peterson et al., 2015; Scheerer et al., 2012). Provided that the current population size remains stable, there seems to be little chance of stochastic extinction of Foskett Dace in the near future.

Our estimates of long-term and contemporary N_e_ suggest that Foskett Spring dace populations have expanded demographically in the last few generations. Long-term N_e_ is defined as the harmonic mean of per-generation N_e_ of a population over evolutionary time (hundreds of generations), whereas contemporary N_e_ measures population sizes within the previous generation or over just a few (Nadachowska-Brzyska et al., 2021). While periodic connectivity with the Coleman Creek population during wet years could help explain the population expansion within Foskett Spring over evolutionary time (see below), the high contemporary N_e_ of Foskett Spring most likely reflects the active restoration work to increase open water habitat at Foskett Spring (U.S. Fish and Wildlife Service, 2019a, 2019b).

Whatever their cause, the high estimates for N_e_ in Foskett Spring help to explain the low inbreeding coefficient (0.036, Table 2). While not zero, the very slight amount of inbreeding certainly represents no cause for concern. This number is nearly zero, and lower than the estimates for the even larger populations in Twentymile and Deep creeks, suggesting that the Foskett Spring population is at least as healthy as those in the stream networks. Indeed, they are arguably faring better than the stream fishes, perhaps because of the lack of any competitors or predators in their habitat. Foskett Dace enjoy sole occupancy of their spring, while dace in Twentymile and Deep creeks share their habitat with several other fishes, including suckers, sculpins, and trouts. The first of these plausibly competes with daces, while the latter two certainly eat them.

As for the refuge population at Dace Spring, the estimated N_e_ of 506 individuals is just a tenth of the value in Foskett Spring, which is perhaps not surprising given that only ∼100 fishes founded the subpopulation at this locality a decade ago. This value of N_e_ is relatively low, and while not at the extremes of 122 individuals that Tian et al. (2022) calculated for *Cyprinodon diabolis*, or the 28 individuals that Black et al. (2017) estimated for *Cyprinodon bovinus*, it may be low enough to incur some risk of stochastic extinction over dozens or hundreds of generations. However, census methods estimated Dace Spring abundance at 1,924 fish (95% CI 1,890–1,965) in 2018, indicating that plenty of recruitment can occur within the spring, and fish in Dace Spring are not substantially more inbred than any other investigated population (Table 2). We conclude there is no immediate risk of losing the refuge population.

Finally, the consistent pattern of reduced observed heterozygosity compared to expected heterozygosity does suggest that allele dropout (due to polymorphisms in restriction enzyme cutsites) may have influenced diversity estimates for all populations (Gautier et al., 2013). If so, the diversity estimates presented herein provide a conservative assessment of population heterozygosity as the true variation could be slightly higher than estimated.

### Recency of origin and intermittent connectivity help to explain the persistence of Foskett Dace

In the introduction, we raised the question of whether luck, genetic diversity, or youth explains the persistence of Foskett Spring Speckled Dace in their tiny habitat. Analyses discussed above indicate that many breeding adults inhabit Foskett Spring, and that these resemble their nearby stream-dwelling cousins in heterozygosity and in their low levels of inbreeding. Those findings indicate that genetic diversity may have played a role in insulating the population from a stochastic drift towards extinction. That said, results from the demographic analysis indicate that genetic diversity does not provide the whole story. Foskett Dace are also much younger and less isolated than previously suspected.

Our demographic model indicates that Western Speckled Dace colonized Foskett Spring from Coleman Creek within the last 500-600 years, not 10,000 years ago during the desiccation of pluvial Lake Warner. That temporal restriction greatly increases the plausibility of the dace’s persistence because it implies that only about 300 generations have passed since isolation, not 5,000. Given that the long-term N_e_ of Foskett Dace is 1,096 (Table 4, Fig. 8), 5,000 generations would have exceeded the number needed for alleles to drift to fixation (4,384, calculated as 4Ne following Kimura & Ohta, 1969), but 300 generations fall far short of that threshold. The reduced duration of persistence also implies far fewer opportunities for rare catastrophic events to cause population crashes. In other words, the danger of drift and the odds of survival are much better over the short term than the long term. Therefore, the need for luck diminishes, and the enigma of persistence largely disappears when the question reframes on the scale of centuries rather than millennia.

The demographic results also suggest that periodic gene flow may have helped to insulate Foskett Dace from the effects of drift (Figs 7, 8). The *D_3_* statistics and demographic models both infer high levels of historic gene flow between Coleman Creek and Foskett Spring after their divergence circa 588 years ago. The *D_3_* statistics also show weak evidence of gene flow between the Coleman Creek and Deep Creek populations. The GADMA model concurred, adding the insight of unequal rates of migration. Gene flow from Deep Creek into Coleman Creek was twice as prevalent as the reverse, and unidirectional gene flow from Deep Creek into Foskett Spring also occurred. Overall, these results help reconstruct how the Foskett Spring population overcame a strong founding bottleneck, in which their originators represented just ∼10% of the ancestral population in the Coleman Valley. Once the spring population established, it grew exponentially (Fig. 8), likely through a combination of recruitment and a periodic influx of migrants from Coleman Creek population whenever it flooded the dry Coleman lakebed. The non-zero levels of gene flow inferred from the Deep Creek population to Foskett Spring (Fig. 8, Table 4) indicate that even dispersal over the rise separating the Coleman subbasin from the remainder of the Warner Valley may have been possible in unusually wet years.

By virtue of these results, the daces of the Warner Valley join a growing list of case studies illustrating how small populations in river systems can persist over evolutionary time by maintaining population sizes and genetic diversity through periodic connectivity (Barson et al., 2009; DeBoo et al., 2019; Mossop et al., 2023; Wilmer et al., 2008; Wong et al., 2004). They reinforce Mossop et al.’s (2015) conclusion that ephemeral waters represent important dispersal routes in arid ecosystems. Such connections, however fleeting, seem to help freshwater fishes in isolated waters to overcome the small population sizes and low genetic diversity (Attard et al., 2022; Black et al., 2021; DeWoody & Avise, 2000; Ward et al., 1994) that could otherwise promote a drift towards imperilment or extinction.

### Reconstructed demography accords with hydrologic history

The patterns of divergence that we reconstruct among the daces in the southern Warner Valley track recent reconstructions of historical hydrography in the northern Great Basin based on pollen records (Minckley et al., 2007) or the prevalence of tui chub bones in Native American middens (Hudson et al., 2021). Following the desiccation of pluvial Lake Warner, warmer and drier conditions persisted in the Northern Great Basin until ca 7,000 years ago (Minckley et al., 2007). During that xeric period, the Coleman subbasin would have remained hydrologically separate from the rest of the Warner Valley, Then, a mesic “Neopluvial” period with higher than present-day precipitation occurred between 5000-3000 years ago (Hudson et al., 2021; Minckley et al., 2007) with the paleobotanical record at the proximate Lily Lake site recording a noted spike in precipitation about 3300 years ago (see Figure 8 of Minckley et al., 2007). High rainfall at these times likely increased the connectivity between the main Warner Valley and the Coleman subbasin. The end of this Neopluvial period coincides closely with our estimated split 3,000-4,000 years ago between the ancestral Deep Creek subpopulation (representing the population in the main Warner Valley) and the Coleman Valley population (Fig. 8). Thus, it sems very likely that a return to xeric conditions three millennia ago isolated the Western Speckled Daces in the Coleman Valley from those elsewhere in the Warner basin.

That subdivision persisted without major change until about 600 years ago when the demographics of both lineages shifted (Fig. 8). In the Coleman subbasin, daces colonized Foskett Spring from Coleman Creek, while in the main Warner basin, the Deep Creek lineage went through a strong bottleneck (Fig. 8). Because F_ST_ values between the Foskett Spring and Coleman Creek populations broadly resemble those between the Deep and Twentymile Creek subpopulations (Table 3), we interpret the bottleneck that GADMA reconstructs in the Deep Creek lineage as representing the subdivision of Twentymile and Deep Creeks, rather than a catastrophic event that eliminated most of the individuals in the population. While we could not test that conjecture directly because GADMA cannot model more than three populations, the isolation of part of the main Warner Valley population in Twentymile creek would certainly reduce the population remaining elsewhere in the valley.

The colonization of Foskett Spring and the presumed split between the Twentymile and Deep Creek subpopulations falls within a period between 1,500 and 300 years ago when precipitation fluctuated around an average slightly higher than present conditions (Minckley et al. 2007). If that reconstruction is correct, then the Coleman and Crump lakebeds would have filled more regularly, allowing daces to access Foskett Spring and creating a lacustrine connection between the Twentymile and Deep Creek stream networks. Those connections would have broken as the most recent rainy period subsided but could have re-established during particularly wet years. Thus, we interpret the low levels of contemporary gene flow that GADMA inferred between Deep Creek, Foskett Spring and Coleman Creek as indicating intermittent and increasingly infrequent watery connections over the last 600 years.

However, it is critical to consider that the timings discussed above depend strongly upon the assumed mutation rate and generation time. GADMA models time in numbers of generations, not in absolute years, and thus the 588-year date of divergence for Foskett Dace (Fig. 8) reflects simple multiplication of 294 inferred generations by two years per generation. That generation time stems from studies of the closely related *Rhinichthys osculus* in streams distant from Oregon (Peden & Hughes, 1981; Wydoski & Whitney, 2003), and it is possible that Foskett Dace mature or reproduce slightly sooner in their predator-free habitat. Similarly, Martin and Höhna (2018) discuss several reasons why small populations of spring dwelling fish might experience mutation rates faster than Recknagel et al. (2013) estimated for cichlids. We used that estimate because we are not aware of any other published estimates for fish mutation rates applicable to SNP datasets like ours but acknowledge that if the true mutation rate is faster, then Foskett Dace might well have colonized their spring even more recently than 600 years ago.

### Historically, what preserved open water habitat in Foskett Spring?

The relatively recent establishment of the Foskett Spring population 600 years ago, rather than thousands of years ago, helps explain the enigma of this small population’s persistence in such a tiny habitat. However, 600 years is still more than enough time for encroaching aquatic vegetation to have rendered the spring uninhabitable. Spring-dwelling fishes often depend upon open water habitat to reproduce (Black et al., 2016; Kodric-Brown, 1977), yet vegetation commonly encroaches spring systems in semi-arid environments without continued disturbance (Kodric-Brown and Brown 2007). Foskett Dace are no exception (Fig. 4) and though they are no longer listed (U.S. Fish and Wildlife Service, 2019a) their population has relied on regular manual removal of vegetation since the fencing of the habitat to prevent livestock access (U.S. Fish and Wildlife Service, 2019b). Cattle were introduced to the northern Great Basin about 150 years ago, leaving a gap of several centuries between the dispersal of dace to Foskett Spring, and the arrival of European settlers and their livestock. What preserved open water habitat in Foskett Spring during this time?

The answer likely lies in the earlier arrival of bison (*Bison bison)* and feral horses, both of which are known to reduce vegetation in Great Basin spring systems (Kodric-Brown & Brown, 2007; Reimondo, 2012). Grayson (2006) reported that bison appeared in the archaeological and paleontological records of south-central Oregon approximately 500 years ago during the cooler temperatures of the Little Ice Age, citing a bison tooth recovered from a dwelling within the Warner Valley dated to between 642 to 257 years ago (Eiselt, 1998; Moore, 1995). Feral horses arrived approximately 300 years ago (Haines, 1938). Though the bison declined to relatively low abundance and were no longer extant by the time of European settlement (Stutte, 2004) the introduction of domestic livestock followed closely behind. Thus, large herbivores have been consistently present on the landscape surrounding Foskett Spring during the entire time that we estimate dace to have inhabited that water body. Indeed, the close match between the arrival of bison and the date that we infer for colonization of the spring raise the intriguing possibility that these large ungulates were key ecosystem engineers that allowed the dace to thrive in a system that vegetation had previously rendered uninhabitable. Prior to modern governmental management, fires and the actions of Native Americans may also have helped to maintain open waters in Foskett Spring (Kodric-Brown & Brown, 2007), though an ethnohistorical study conducted slightly to the north of this region concluded that human-set fires were much more prevalent in mixed conifer zones than in xeric shrub grasslands like those that surround Foskett Spring (Steen-Adams et al., 2019).

### Conclusions and future directions

This study demonstrates how modern demographic analyses, rich genetic datasets and improved geographic sampling can refine and rewrite our understanding of biogeography, even in well-studied systems. By including specimens from a previously undiscovered population and by coaxing dates of divergence among populations from a large panel of SNPs, we revealed that the riddle of Foskett Dace’s long persistence stemmed from an inaccurate assumption about their age and from the partitioning of knowledge about local fauna and hydrology along state lines. Though many biologists have studied Foskett Dace before (including two authors of this contribution), all had overlooked the Nevada population that provided the key to understanding the origins of one of Oregon’s endemic fishes. So, our results illustrate the importance of questioning one’s assumptions, and how revisiting old questions with new tools and fresh eyes can yield unexpected insights.

Foskett Dace represent just one of the many locally endemic aquatic species occurring in deserts around the world, and many of these have inspired or required intense conservation efforts. Given how much our understanding of the historical demography and biogeography of Foskett Dace shifted because of this study, it might prove informative to reevaluate our knowledge of other desert species. Are some older or younger than commonly believed? Have we missed other components of diversity in the deserts of our planet? Such studies could hold particular value in taxa for which no recent genetic studies have been performed, or where no comprehensive surveys have searched for undiscovered populations in forgotten bodies of water.

As for Foskett Dace themselves, our results reveal their genetic health and support a prediction of their continued persistence. Genetically, they are surprisingly diverse, and appear to thrive within their spring and outflow stream. Given the existence of the refuge at Dace Spring and the regular attention that their managers place on ensuring that enough open water exists to support recruitment, their future seems secure for now. That said, monitoring should continue, and periodic estimation of heterozygosity and effective population size would help protect this isolated and potentially fragile population. Fortunately, we know now of a population of close relatives in Coleman Creek with which genes have flowed in the last several centuries. If monitoring ever reveals declining diversity in Foskett Spring, the Coleman Creek population provides an obvious reservoir of additional diversity that could be used in genetic rescue.

Finally, our results do not speak to the origins of the unusual morphology of Foskett Dace, which possess a shorter lateral line and more posterior dorsal fin than other daces in the Warner Valley. These features led Carl Bond to originally propose subspecies status for this isolated population. Hoekzema (2013) confirmed the morphometric diagnosability of Foskett Dace and demonstrated that they have, on average, eight fewer pored lateral line scales than do stream-dwelling daces in the main Warner Valley. However, that study did not include Coleman Creek dace, because no one knew that they existed. Do Coleman Creek dace more closely resemble Foskett Dace or Deep Creek dace in morphology? Does the morphometric variation result from phenotypic plasticity, or from local adaptation? If the latter, then the spring phenotype has arisen in no more than 300 generations, which is relatively rapid in evolutionary terms, though not unreasonably so (Willoughby et al., 2018). If these morphologies do adapt Foskett Dace to their unusual environment, then those adaptations might also help explain their persistence (Attard et al., 2022). The presence or absence of adaptive variation would also help reveal whether Foskett Dace have evolved substantially since their isolation. That determination could in turn inform any future debate about whether Foskett Dace qualify as a distinct population segment, an evolutionarily significant unit, or neither (Pennock & Dimmick, 1997). Answers to these questions will require morphological comparisons between Foskett Dace, Coleman Dace and dace from elsewhere in the Warner Valley, a screen of the SNP panel for loci under selection, and construction of a reference genome for *Rhinichthys osculus.* When complete, that work will likely reveal that this small fish in an isolated spring has not yet yielded all its secrets.

## Data Availability Statement

Unprocessed sequence data has been deposited to NCBIs Sequence Read Archive under Bioproject PRJNA934901. All scripts and barcode files have been deposited to github (https://github.com/Andrew-N-Black/Speckled_dace).

## Acknowledgments

We thank Jimmy Leal, Grace Haskins, and the Bureau of Land Management for funding this project under cooperative agreement #L20AC00103. Justin Miles (ODFW) provided logistical support during the field collections. Michele Weaver (ODFW) helped to secure the Oregon collection permit, and Travis Hawks (NDOW) researched the management history of Coleman Creek at our request. Katie Carter, Mark Dasenko, and Liz Zepeda at OSU’s Center for Quantitative Life Sciences performed the lab work needed to generate raw data from the tissue subsamples. We also thank Kendra Hoekzema for sharing her thoughts about Western Speckled Dace diversity in the Great Basin, Anna Brüniche-Olsen for providing advice on demographic modeling, the DeWoody lab for providing useful feedback on earlier versions of this manuscript, and Adam Hudson for insights on the paleohydrology of the northern Great Basin.

## Author Contributions

BLS and AB conceived the study and secured funding. BLS, HA and FM conducted fieldwork. BLS and HA prepared samples for sequencing. AB and SM analyzed the data. All authors prepared figures. BLS led the writing and editing, with SM, AB and FM contributing sections of text. All authors revised the manuscript and approved its submission.

